# Tuning viscoelasticity and fine structure of living materials via synthetic adhesion logic and rheological perturbations

**DOI:** 10.1101/2025.06.04.657808

**Authors:** Stefana A. Costan, Kira Hallerbach, Samuel Y. Kim, Chris Camp, Minkyu Kim, Ingmar H. Riedel-Kruse

## Abstract

Engineered living materials (ELMs) at the multicelluar level represent an innovation that promises programmable properties for biomedical, environmental, and consumer applications. However, the rational tuning of the mechanical properties of such ELMs from first principles remains a challenge. Here we use synthetic cell-cell adhesins to systematically characterize how rheological and viscoelastic properties of multicellular materials made from living bacteria can be tuned via adhesin strength, cell size and shape, and adhesion logic. We confirmed that the previous results obtained for non-living materials also apply to bacterial ELMs. Additionally, the incorporation of synthetic adhesins, combined with the adaptability of bacterial cells in modifying various cellular parameters, now enables novel and precise control over material properties. Furthermore, we demonstrate that rheology is a powerful tool for actively shaping the microscopic structure of ELMs, enabling control over cell aggregation and particle rearrangement, a key feature for complex material design. These results deepen our understanding of tuning the viscoelastic properties and fine structure of ELMs for applications like bioprinting and microbial consortia design including natural systems.

## Introduction

A central goal of engineered living materials (ELMs) field is to utilize genetic engineering techniques to create novel and functional materials with programmable properties by manipulating biological systems [1]. Multicellular (MC) living systems are an important class of ELMs [2, 3] that are inspired by prokaryotic biofilms and eukaryotic organisms and which display a large variety of morphologies (3D structures) and patterns (cell types) [4, 5, 6, 7, 8, 9]. The control over cell-cell adhesion is key to control structural integrity, morphology, patterning and functionality of multicellular ELMs. The molecular and cellular biophysical properties of a bacterial synthetic adhesin toolbox [10] were quantitatively characterized recently and which then allowed the bottom-up prediction and tuning of the tensile strength of such ELMs [11]. Other synthetic adhesin toolboxes and their applications have been reported [12, 13]. Synthetic biology aims to e!ciently design and construct complex living systems from standardized and thoroughly characterized components [14, 15]. This also promises the creation of innovative smart materials featuring diverse length scales, design strategies, and matrix types for biomedical, environmental, and consumer applications [16] such as bioprinting sensors [17], therapeutic patches [18], in vivo drug-delivery vehicles targeting gastrointestinal microbiota or tumor cells [19, 20], and microbial consortia for distributed biochemical synthesis [21].

Bacterial rheological studies have been conducted for cultures, biofilms, and scaffold-embedded cells, and the effects of various parameters such as cellular components [22, 23, 24, 11], external stimuli [25], or non-living components [26, 27] have been previously investigated. ELMs made entirely of bacterial cells lack systematic viscoelastic characterization [28]. Synthetic adhesins provide many opportunities to fine-tune properties from the molecular to the macroscopic material level, such as modulating the number of adhesin pairs between cells [29, 10], modulating the linker length between membrane and adhesin domain [30], utilizing different cell shapes [10], and combinatorial use of an adhesin library, often referred to as combinatorial adhesin logic or, loosely interchangeably, adhesion code [10, 6, 31] (Fig. 1a). The modeling of MC ELMs is currently in its nascent stages, with researchers exploring diverse approaches to understand and design these complex systems [32, 16]. Therefore, such systematic characterizations can significantly contribute to the development of these algorithms. Rheological measurements, a cornerstone of material characterization, offer a systematic approach to understanding viscoelastic properties [33, 34] (Fig. 1b). By using techniques such as flow sweeps to measure viscosity [35] (Fig. 1c) and amplitude sweeps to determine storage and loss moduli [36] (Fig. 1d), researchers can quantify material rigidity and fluidity. Although widely applied to non-living suspensions [37, 38, 39, 40], these methods remain underexplored for ELMs composed solely of bacterial cells [41], reflecting a wider growing interest in active colloid suspensions [42].

**Fig. 1:**
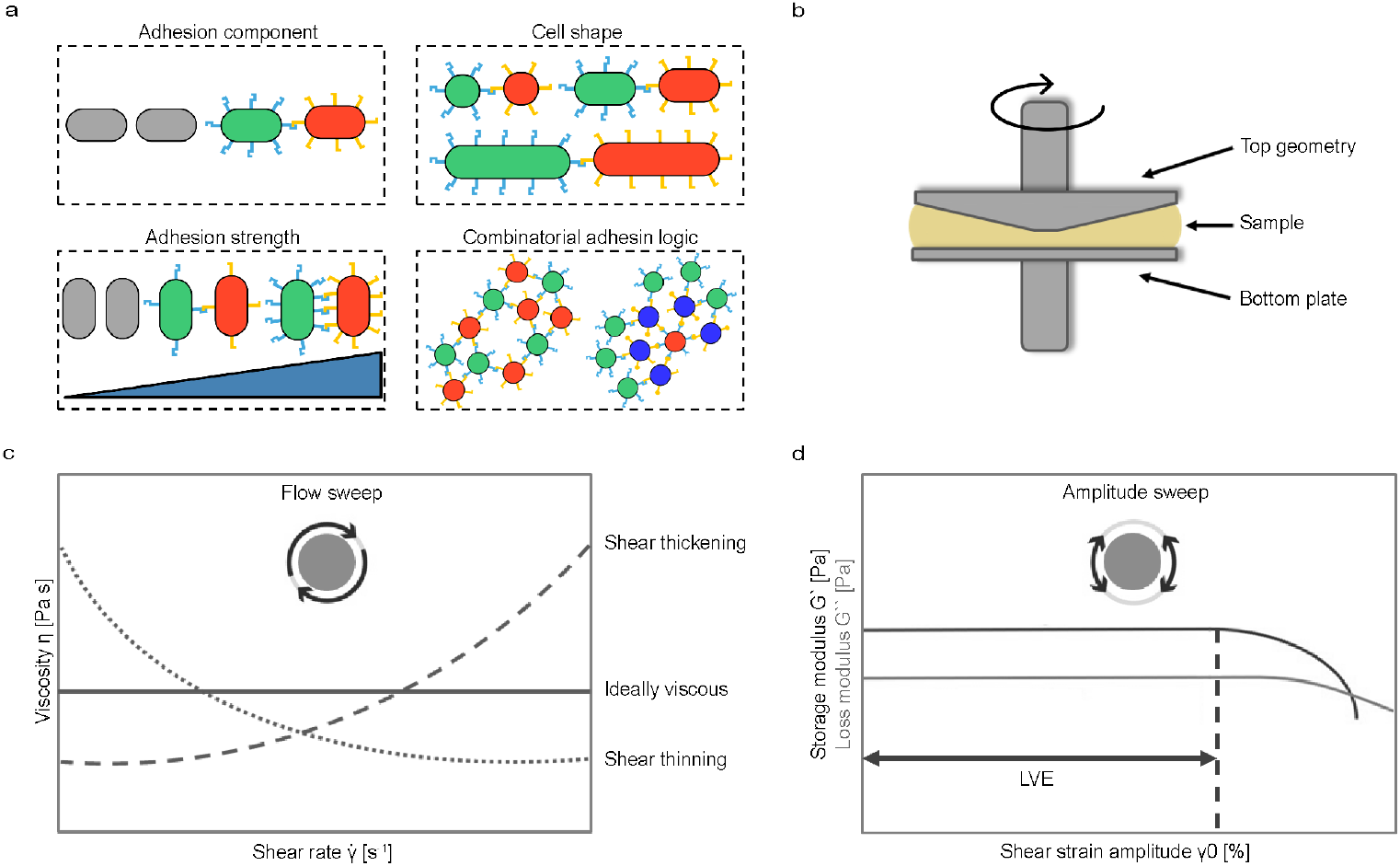
Key properties of engineered living material (ELMs) can be tuned via synthetic adhesins and other cellular parameters. **a** Cellular parameters for heterophilic adhesin constructs that can be varied for rheology measurements: integration of the adhesion component, adhesion strength, cell shape, combinatorial adhesion logic. **b** Illustration of the rheometer and its main components, i.e., the fixed bottom plate and the top conical geometry, between which the sample is placed. **c** Examples of typical viscosity curves that can be measured via rheology. **d** Examples of elastic modulus curves that can be measured via rheology, where LVE represents the linear viscoelastic range.

In this paper, we used rheology to systematically characterize how synthetic adhesins allow the tuning of the viscoelastic and structural properties of ELMs. We varied parameters of adhesion strength, logic and cell shape (Fig.1a) in a methodical fashion to then measure the viscoelastic response and characterize the behaviour of these ELMs. We verified if the rheology trends observed for non-living materials [37, 38, 39, 40] hold true for bacterial ELMs, and we evaluated the effect that the addition of the adhesion component has on material properties. Based on previous findings of particle rearrangement inside the material due to increased shear rate [39], we concluded with a demonstration on how rheology can be used as a tool for tuning the miscroscopic material structure, i.e., the mixing, demixing and generation of particle aggregates of different types and sizes.

## Results

### Integration of synthetic adhesion (dual-component) in ELMs

We first investigated how the expression of synthetic adhesins on the outer membrane of bacterial cells affects the material properties of ELMs. Adhesion was promoted by adding 100 ng/mL anhydrous tetracycline (aTc) to overnight cultures of heterophilic mixtures of nanobody (Nb3)-expressing and antigen (Ag3)-expressing cell strains, as previously described [43, 11] (Fig. 2a top). To confirm adhesin expression, we used confocal imaging to compare these heterophilic mixtures which were also transformed to express cytoplasmic fluorescence, with wild-type (wt) cells (Fig. 2a bottom). Upon measuring the viscosity of the cell pellets formed from these cell cultures, we observed that wt cells (*n*=7 repeats across 4 days) exhibited higher viscosity than adhesin-expressing cells (*n*=9 repeats across 5 days) at low shear rates (Fig. 2b) (Student’s t-test, p = 0.0019). This result may be attributed to differences in cell clustering before centrifugation: confocal images showed that adhesin-expressing cell clusters had gaps between cells, leading to less densely packed pellets compared to wt cells. At higher shear rates the viscosity difference was reversed, but not significantly (Student’s t-test, p = 0.22).

**Fig. 2:**
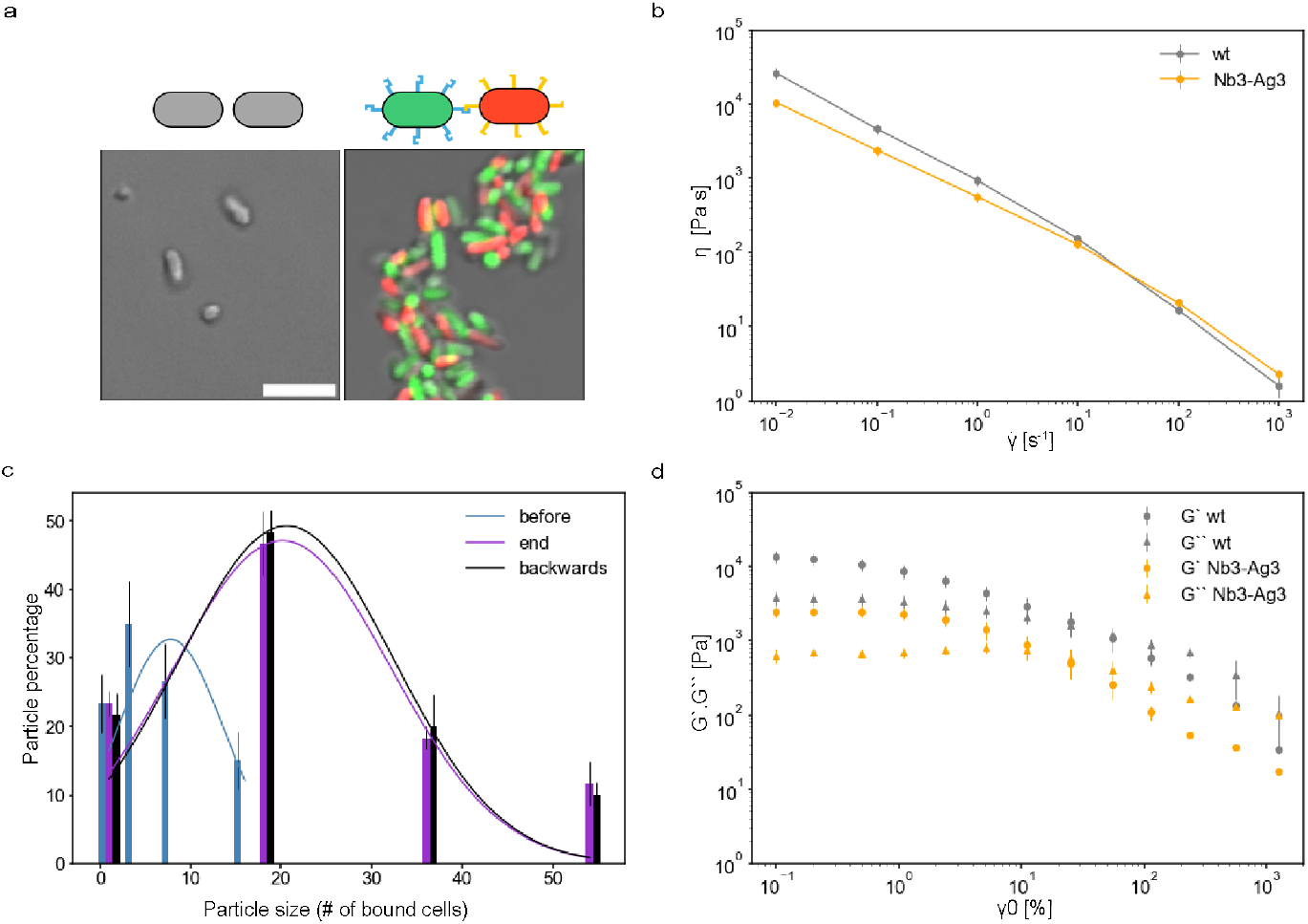
The integration of the adhesion component changes the rheological properties of the ELM. **a** Illustration of the bacterial cells with and without adhesins (top) and confocal images of the wild type (wt) cells and the induced heterophilic mixture of the Nb3-Ag3 adhesin pair in solution before centrifugation (bottom). Scale bar = 5 μm. **b** Measured viscosity for the Nb3-Ag3 and the wt cell pellet. **c** Cluster size distribution for the Nb3-Ag3 material dissolved in solution before viscosity measurement, at the end of the measurement and after a second round with the shear rate order reversed (backwards measurement). **d** Storage and loss modulus of the wt and the Nb3-Ag3 cell pellet. Data presented as mean *±* sem unless noted otherwise.

To understand how rheology affects the internal structure of the material, we examined the cluster size distribution in the ELM made from induced heterophilic cells at different stages: before any rheology measurements (blue), at the end of one complete round of viscosity measurement (purple) and after a second round with the shear rate order reversed (backwards measurement, black) (Fig. 2c). We found that after one round of rheology measurements the clusters size increased by a factor of 3 from the initial size with the mean value for the number of cells bound changing from 7.8 ± 2.4 (blue) to 20.1 ± 4.1 (purple) (mean ± sem throughout text, unless noted otherwise. Student’s t-test, p= 0.02, *n*=9 repeats across 5 days). There was no significant difference in cluster size distribution between after the first measurement (purple) and after the second measurement with reversed shear rate order (black) (Student’s t-test, p ≥ 0.1).

We conducted oscillation tests to determine the viscoelastic properties of the cell pellet, varying the shear strain amplitude between 10^−1^ and 10^3^% (Fig. 2d) (*n* = 3 repeats across 2 days). The linear viscoelastic (LVE) region (where the material does not get degraded by the strain amplitude) for the pellet from adhesin-expressing cells was between 0.1 and 5% strain. In contrast, the wt cell pellet had a shorter LVE region, indicating greater brittleness (Fig. 2d), which aligns with previous tensile strength experiments [11]. As a control, we compared rheological data for the wt and uninduced heterophilic cells, finding no significant differences; thus, we combined the data for both viscosity and viscoelasticity analyses (Supplementary Note 1).

### Material tuning via the adhesin strength

We next examined how the viscosity of ELMs varies with the level of adhesin induction. By imaging heterophilic Nb3-Ag3 cell mixtures in solution before centrifugation, we found that cluster size increases proportionally with the induction level (Supplementary Note 2). We measured the viscosity for ELMs induced with aTc concentrations ranging from 0 to 300 ng/mL (Supplementary Note 2). At a shear rate of 310 s^−1^, we observed a gradual increase in viscosity with increasing induction level, prompting us to use this shear rate for further analysis. The viscosity data were fitted to a Hill equation (Fig. 3a):

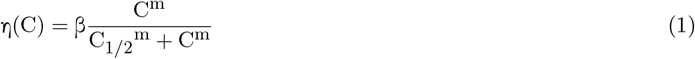

where, C is the inducer concentration, C_1/2_ being the inducer concentration yielding half of the maximum viscosity and η (C) the viscosity value as a function of inducer concentration, m is the Hill coe!cient and β is the maximal expression level of the promoter. From the fit, we determined C_1/2,conc_ to be 79 *±* 19 ng/mL aTc and the Hill coe!cient m to be 1.7 *±* 0.6 (*n* = 4 repeats across 2 days), which are consistent with those reported in the previous study [11] (Student’s t-test, p ≥ 0.1).

**Fig. 3:**
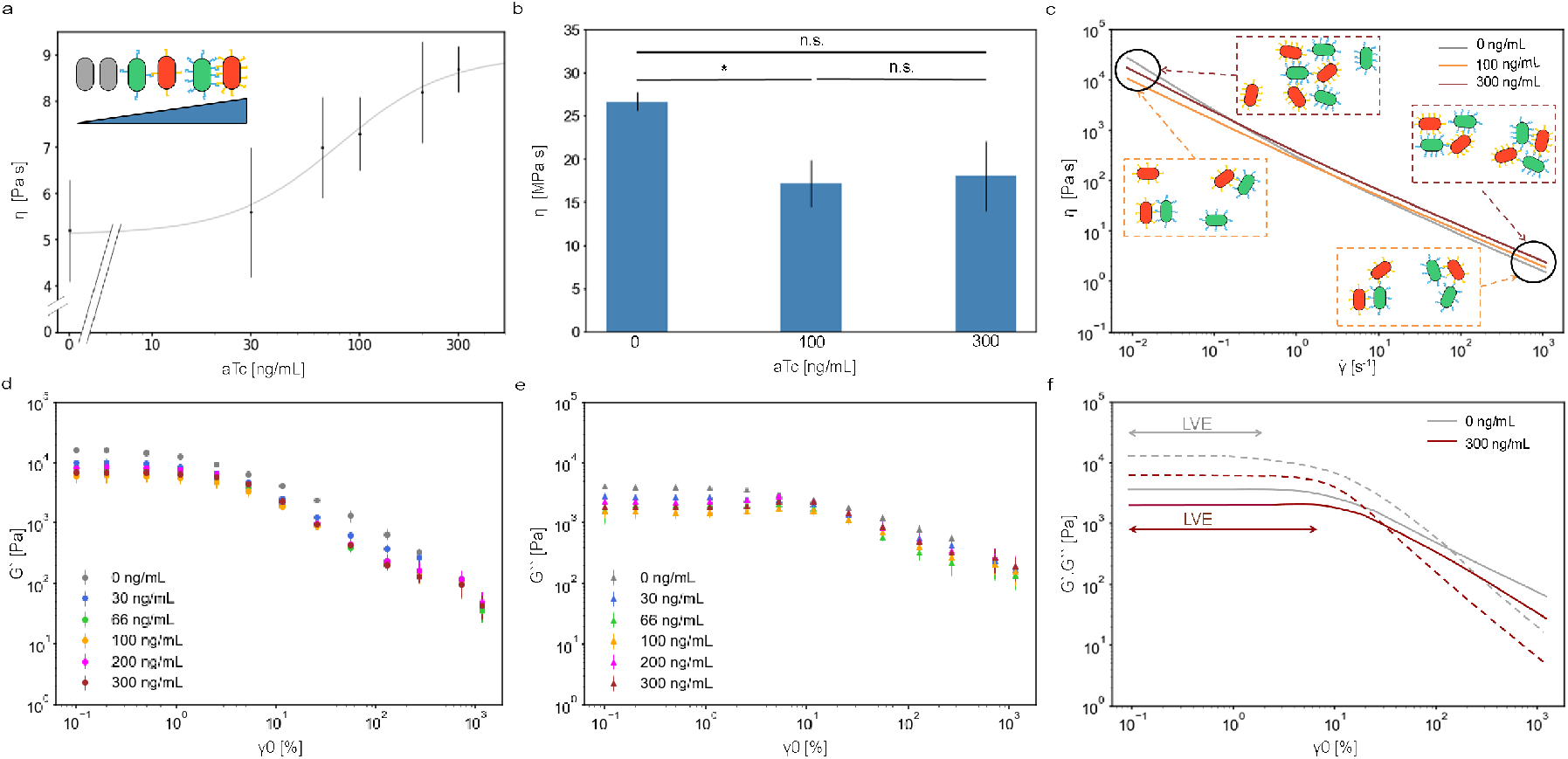
The rheological properties of the ELM can be tuned by the adhesin number. **a** Viscosity measured at a constant shear rate of 310 s–1, across a range of aTc concentrations between 0 to 300 ng/mL for a heterophilic mixture of the Nb3-Ag3 adhesin pair (the same as above). **b** Comparison between viscosities at a low shear rate of 0.1 s–1 for the uninduced, intermediary and maximally induced cells. **c** Qualitative viscosity model for di!erent induction levels and a schematic of their associated cluster sizes changing with an increasing shear rate; top insets for 300 ng/mL aTc and bottom insets for 100 ng/mL aTc; 0 ng/mL induction did not exhibit any di!erence in the confocal images for the before and after rheology measurements. **d** Storage modulus measured for materials induced with aTc concentrations between 0 and 300 ng/mL aTc. **e** Loss modulus measured for materials induced with aTc concentrations between 0 and 300 ng/mL aTc. **f** Qualitative model for the e!ects of adhesion strength on the viscoelasticity of the material, where solid and dotted lines represent the storage and loss modulus, respectively.

We then investigated the behavior of the materials at lower shear rates (0.1 s^−1^). At this shear rate, the viscosity of uninduced cells (0 ng/mL aTc) was statistically similar to that of maximally induced cells (300 ng/mL aTc) (Student’s t-test, p = 0.09, *n* = 4 repeats across 2 days). Cells induced with an intermediary concentration of aTc (100 ng / ml aTc) exhibited a significant lower viscosity compared to uninduced ones (Student’s t-test, p = 0.002), but not significantly different from maximally induced cells (Student’s t-test, p = 0.86) (Fig. 3b). To understand the structural basis for these viscosity differences, we examined the material microscopic structure before rheology measurements, as this structure corresponds to the state during low shear rate viscosity tests. Samples were dissolved in PBS and imaged. At 0 ng/mL aTc induction the material consisted of monodisperse single cells (Supplementary Note 2). At intermediate induction levels we observed polydisperse clusters averaging 7.8 *±* 2.4 cells (Fig. 2c) (Supplementary Note 2). At maximal induction level, large clusters of 101 *±* 35 cells were present (Supplementary Note 2). These observations led us to propose a qualitative model for ELMs with varying particle sizes, influenced by induction level, which parallels models for non-living materials [44, 37, 38].

We performed small amplitude oscillatory shear rheology to characterize the viscoelastic properties of the materials across different induction levels, measuring the storage modulus (G′) and loss modulus (G″). At lower shear rates, G′ exceeded G″, indicating solid-like behavior, while at higher shear rates G″ dominated, signifying liquid-like behavior (Supplementary Note 2). For induced cells, the LVE and crossover point were consistent, with an yield stress (σ_y_) of 55 *±* 11 Pa and a flow stress (σ_f_) of 395 *±* 53 Pa, respectively (*n* = 3 repeats across 2 days). In contrast, uninduced cells exhibited a shorter LVE and a higher crossover point with σ_y_ of 120 *±* 25 Pa and σ_f_ of 933 *±* 229 Pa, respectively. This suggested that uninduced ELMs may be more prone to brittle fracturing compared to induced ones. Finally, we developed a qualitative model to describe the viscoelastic behavior of these ELMs, illustrating the shorter LVE and a higher crossover point for cells lacking adhesins compared to induced ones (Fig. 3f).

## Material tuning via the cell shape

We next investigated how cell shape influences the properties of ELMs. To modulate the cell aspect ratio, we used spherical cell strains from our adhesin library [10] or treated cells with cephalexin to induce elongation [45] (Fig. 4a). We imaged the cells under two conditions: no induction (Fig. 4a top) and 100 ng/mL aTc (Fig. 4a bottom). Initially we measured the viscosity of ELMs composed of uninduced cell mixtures. At low shear rates, we observed a trend of increasing viscosity with cell aspect ratio, although this was not significant (Student’s t-test, p ≥ 0.2) (*n* = 4 repeats across 2 days) (Fig. 4b). At higher shear rates, the viscosities were comparable across different aspect raios (Student’s t-test, p ≥ 0.3). For induced cells, viscosity measurements revealed no significant differences at either low or high shear rates across varying aspect ratios (Student’s t-test, p ≥ 0.3) (Fig. 4c).

**Fig. 4:**
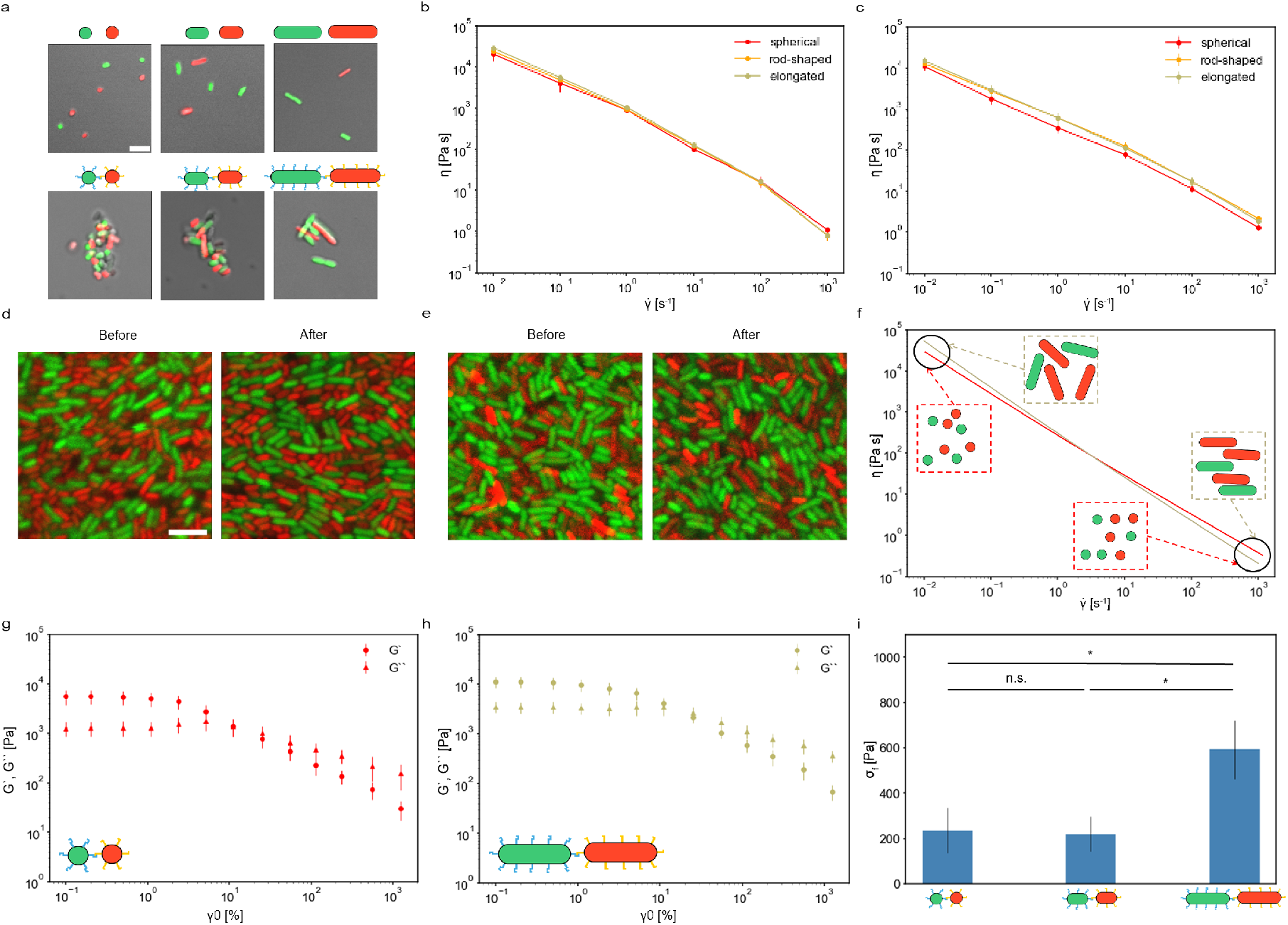
Cell shape and the adhesion component regulate the rheological properties of the ELM. **a** Confocal imaging of the uninduced (top) and induced (bottom) heterophilic Nb3-Ag3 cell mixtures of di!erent aspect ratios (the ratio between length and diameter of 1, 2 and 4 from left to right, respectively) while in solution, before centrifugation. Scale bar 5 μm. **b** Measured viscosity of the uninduced cell pellets consisting of di!erent cell shapes. **c** Measured viscosity of the induced cell pellets consisting of di!erent cell shapes. **d** Confocal images of the uninduced cell pellet before and after rheology. Scale bar in d 5 μm. **e** Confocal images of the induced cell pellet before and after rheology. Scale bar as in d. **f** Qualitative model for the e!ects of cell shape on the material viscosity when cells are uninduced based on the (not yet significant) data trends and literature expectations; inset schematics show the elongated (top) and spherical (bottom) cells orienting with the flow. **g**,**h** Measured storage and loss modulus for the material consisting of cells with di!erent aspect ratios. **i** Comparison between flow stresses as a function of the cell aspect ratio.

To asses cell alignment induced by rheology, as suggested by prior studies on non-living suspensions [40], we imaged the elongated cell pellets before and after rheology measurements (Fig. 4d,e). We quantified cell orientation relative to an arbitrary axis by calculating the circular mean of angles from 20 cells (*n* = 4 confocal images across 2 day). For uninduced cells the circular mean increased significantly from Z_no,bef_ = 0.09 *±* 0.04 to Z_no,aft_ = 0.28 *±* 0.04 (Student’s t-test, p = 0.002). In contrast, for induced cells the circular mean only showed a non-significant increase, from Z_ind,bef_ = 0.11 *±* 0.04 to Z_ind,aft_ = 0.16 *±* 0.05 (Student’s t-test, p = 0.9). This suggests that the lack of adhesins allowed the elongated cells to align more readily when the shear strain was applied by the rheometer. Based on these (not yet significant) trends in the data as well as expectations from the literature [40, 39], we developed a qualitative model illustrating influence of cell shape on ELM viscosity (Fig. 4f).

We conducted oscillatory tests on the cell pellets to further characterize their viscoelastic properties (Fig. 4g,h, Supplementary Note 3). Analysis of the data revealed that elongated cells exhibited higher G_′_ and G″ compared to spherical cells (Student’s t-test, p *<* 0.05) (Fig. 4g,h), which is in agreement with studies done on non-living suspension and is explained by elongated rod particles being able to enhance G′ due to increased particle-particle interactions [46]. Moreover, higher aspect ratios increased the storage modulus by promoting particle orientation along the flow direction, leading to a more elastic response [47]. Similarly, we determined an increase in the yield stress, σ_f_, (Student’s t-test, p = 0.04) with increasing cell aspect ratio (Fig. 4i), explained by the ability of elongated particles to form networks and lead to stronger gel structures, requiring greater stress to initiate flow [48].

### Material tuning via combinatorial adhesin logic and rheological perturbations

Lastly, we investigated how multiple rheology iterations affect ELMs with or without adhesins, building on our earlier observation of cell rearrangement under shear stress (Fig. 4d). Equipped with a combinatorial cell–cell adhesion logic [10], we investigated dual-component materials of Nb3-Ag3 cells (Fig. 5a top) as well as tri-component materials of Nb2, Ag2/Ag3, and Nb3 cells (Fig. 5a bottom) and different mixing ratios. First, we established a baseline using a two-component mixture of 1:1 spherical heterophilic Nb3 and Ag3 cells (Fig. 5a top). Qualitative comparison of confocal images suggested that after rheology induced cells became more uniformly distributed, indicating enhanced mixing on a larger scale (Fig 5b), whereas for uninduced cells we did not see any rearrangement. This suggested that adhesion plays a crucial role in mediating structural changes during rheology, and we therefore hypothesized that a higher level adhesion (tri-component) would promote prominent quantitative rearrangement of cells inside a material.

**Fig. 5:**
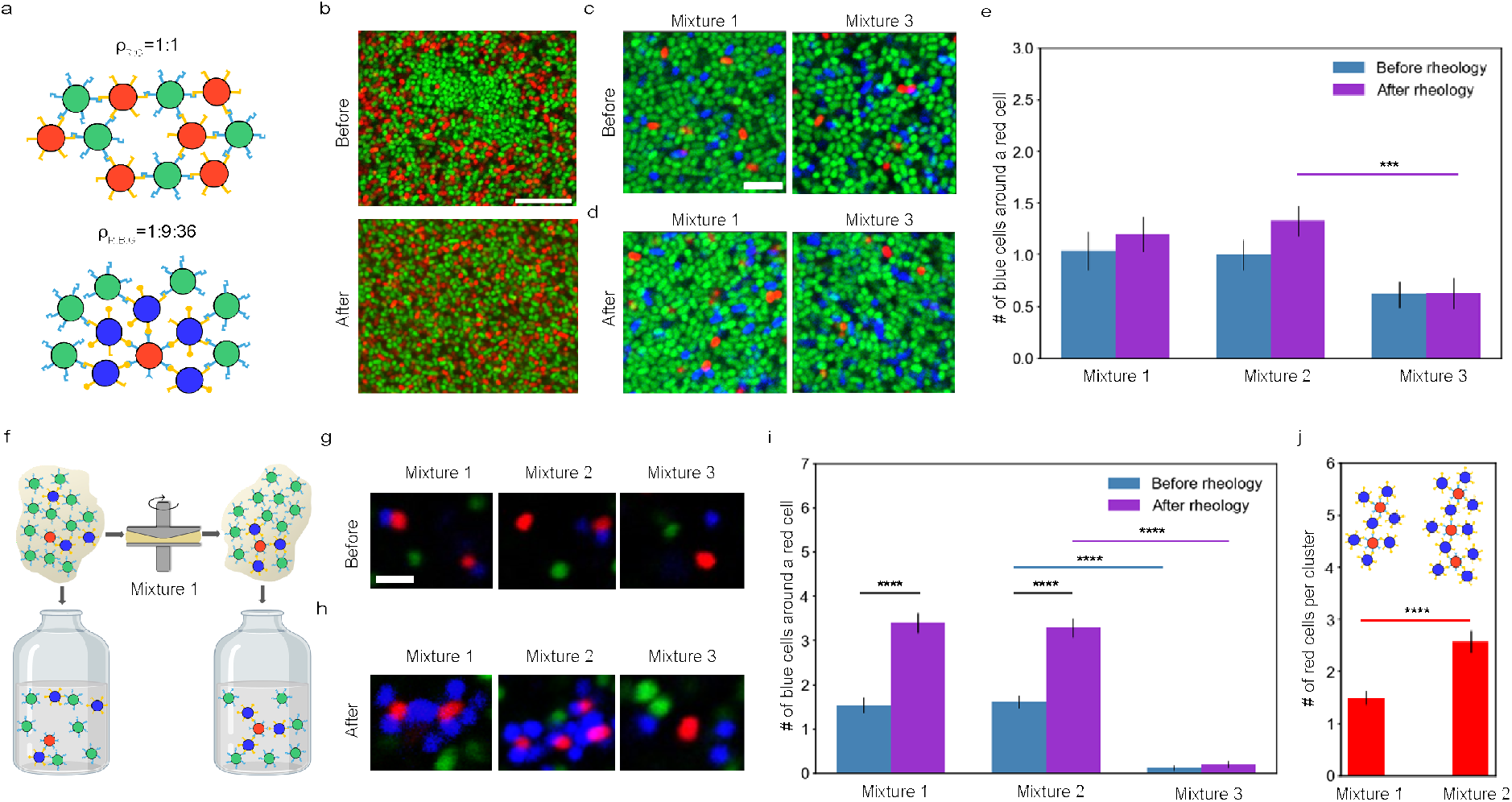
Adhesin-expressing cells rearrange inside the ELM due to multiple rheology iterations. **a** Schematic showing the combinatorial adhesin logic for the formation of clusters in b-j; dual-component (top) and tri-component (bottom) materials, as well as the density ratios β between green (G), red (R), and blue (B) cells. Figure adapted from [10]. **b** Mixing and homogenozation phenomenon inside a 2-strain mixture (1:1) of heterophilic Nb3-Ag3 cells, where a more homogeneous mixture is obtained after rheology on larger scale, illustrated through confocal images of the cell pellet before and after rheology testing. Scale bar 10 μm. **c**,**d** Demixing / sorting phenomenon inside a heterophilic 3 strain mixture of 1:9:36 (R:B:G) observed through confocal images of the cell pellet before and after rheology testing, respectively. The mixtures characteristics are a follows: in mixture 1 the adhesins were induced in all cells; in mixture 2 the adhesins were only induced in Nb2 (red) and Ag2/Ag3 (blue); and in mixture 3 no adhesins were induced in any cell. Scale bar 5 μm. **e** Number of blue cells surrounding the red ones inside the cell pellet before and after rheology. **f** Illustration showing the methodology and expected e!ects of rheology on the internal structure of the material in both the pellet and resuspended form. Note: schematic is for illustrative purposes only. We did not observe a clear rearrangement of the cells in pellet form due to limitations in the 3D image reconstruction. **g**,**h** Cluster comparison between the three mixtures before and after rheology and resuspension in PBS, respectively. Scale bar 2 μm. **i** Number of blue cells surrounding the red ones in the cell suspension from before and after rheology measurements. **j** Comparison for the number of red cell per cluster in the cell suspension after rheology between Mixture 1 and 2. Note: green cells not included for simplification of inset schematic.

Next, we employed a tri-component system consisting of Nb2, Ag2/Ag3, and Nb3 cells, using a sequential mixing protocol to generate layered clusters as previously described [10]. Specifically, we mixed spherical Nb2 (red) and Ag2/Ag3 (blue) cells in a ratio of 1 to 9, incubated for 1 h, then added Nb3 (green) cells at a 36:1 ratio relative to the Nb2 cells and incubated for another hour (Fig. 5a bottom). We investigated mixtures with different combinations of induction: Mixture 1: adhesins were induced in all cells; Mixture 2: adhesins were only induced in Nb2 (red) and Ag2/Ag3 (blue); and Mixture 3: no adhesins were induced in any cell. After centrifugation, the pellets were imaged in slide chambers with a height of 0.3 mm to assess cell distributions before (Fig. 5c) and after (Fig. 5d) rheology. For Mixtures 1 and 2, we observed a slight yet not statistically significant increase in the association of blue Ag2/Ag3 cells with red Nb2 cells after rheology: from 1.04 ± 0.19 to 1.21 ± 0.17 for Mixture 1 (p = 0.5) and from 1.00 ± 0.16 to 1.33 ± 0.15 for Mixture 2 (p = 0.1), suggesting a tendency towards greater clustering (n = 6 randomly picked cells for each confocal slice, with 2 slices for each one of the 2 stacks, at 0 and 10 σm) (Fig. 5c-e). In contrast, Mixture 3 showed no significant change (1.00 *±* 0.17 and 1.08 *±* 0.19 for before and after rheology, respectively) (Fig. 5c-e). With further analysis we controlled that these effects are not due to irregular mixing ratio of these mixtures (data not shown). Hence, this data suggested that mechanical perturbations applied through multiple rheology iterations did lead to structural sorting in the presence of cell-cell adhesins, but not in the case of non-adhering cells.

To test, whether these rearrangements effects were real and what their microscopic impact was, we performed re-suspension experiments: we dissolved the cell pellet in a solution of PBS before and after rheology measurements (Fig. 5f), and we also imaged the cells before and after (Fig. 5g,h). We found that for Mixtures 1 and 2, the number of Ag2/Ag3 (blue) cells surrounding each Nb2 (red) cell increased significantly after rheology, from 1.58 *±* 0.11 to 3.35 *±* 0.15 (p ≤ 0.0001; Fig. 5i). Therefore, we conclude that rheology promoted cell rearrangement and that persisted even after resuspension.

While we previously established that rheology increased red-blue binding (Fig.5g-i), we were finally interested to see if it also increased the size of the red-blue clusters (as measured by number of red “center” cells in a cluster) in conjunction with the adhesins expressed on the excess cell type (in this case Nb3 green). We compared Mixtures 1 and 2, noting that in Mixture 2, where Nb3 cells were uninduced and acted as fillers, larger clusters formed post-rheology compared to Mixture 1 where Nb3 cells were induced, with cluster sizes containing 2.58 ± 0.22 red cells versus 1.50 ± 0.14, respectively (Fig 5j). This parallels non-living concentrated suspensions where smaller particles facilitate larger aggregate formation [49]. To confirm this, we performed experiments without Nb3 filler cells, which showed no increase in cluster size after rheology (Supplementary Note 5). Thus, multiple rheology iterations led to significant cell rearrangement in ELMs, particularly when adhesins were expressed, offering a method to engineer complex material structures.

## Discussion

In summary, we presented a systematic investigation into the rheological properties of ELMs composed of bacterial cells, highlighting the key role of synthetic cell-cell adhesins in tuning these properties. We studied how the rheological ELM properties can be quantitatively tuned by key parameters such as the expression of cell-cell adhesins, the level of induction, cell shape, and the mixing ratios of multi-component adhesin constructs (Fig. 1). Our findings confirmed that fundamental rheological principles established for non-living materials also applied to bacterial ELMs [37] -[40]. However, the incorporation of synthetic adhesins, combined with the adaptability of bacterial cells, enabled precise control over material properties through a systematic investigation of multiple parameters, each explored through individual variation, marking the first instance of such a comprehensive approach in the literature [16, 22, 50, 51]. This study not only elucidated the specific contributions of each parameter but also equipped us with a diverse set of tools and multiple venues for living material optimization.

Specifically, we observed that the expression of synthetic adhesins significantly influenced the viscosity of ELMs (Figs. 2,3). Wt cells, lacking adhesins, exhibited higher viscosity at low shear rates compared to adhesin-expressing cells. This can be explained with previous findings on concentrated particle suspensions [37], where few larger objects have a lower viscosity than many smaller objects, and where the synthetic adhesins now lead to aggregation of cells into clusters, thereby reducing the overall number of “particles” in solution (given the same cell density). This difference diminished at higher shear rates due to particle rearrangement with respect to the flow direction, also in agreement with established suspension rheology [37] (Fig. 2). The level of adhesin induction also played a crucial role, with intermediary induction levels leading to polydisperse solutions where smaller particles acted as lubricants, reducing viscosity [44] (Figs. 2, 3 and Supplementary Note 2). Our oscillation testing revealed that wt cells have a shorter LVE region compared to adhesin-expressing ELMs, indicating greater brittleness, consistent with previous tensile strength experiments [11].

Cell shape was another key parameter affecting material properties (Fig. 4). Uninduced elongated cells showed higher viscosity at low shear rates compared to spherical cells, but this effect reversed at higher shear rates, likely due to the ability of uninduced cells to align with the flow [52, 39]. When adhesins were present, cells bound together, limiting their ability to reorient. Furthermore, our data showed that cell shape influenced not only viscosity but also viscoelastic moduli, with elongated cells exhibiting higher storage and loss moduli and a higher flow stress, suggesting enhanced structural integrity under shear (Fig. 4).

This work also provides the first demonstration that the rheology in combination with synthetic adhesins can be used to actively shape the microscopic structures of ELMs. By applying controlled shear forces, we induced mixing, demixing, and the formation of specific cell patterns (Fig. 5). For instance, multi-component adhesin systems combined with multiple rheology iterations enabled the formation of well-defined clusters, showcasing the potential to engineer spatially diverse microbial consortia within ELMs [32]. This capability is especially significant for applications requiring precise control over material architecture, such as bioprinting [17] or the creation of spatially defined consortia for drug synthesis [53], as well as understanding the microscopic changes that occur when different flow conditions induce different morphologies in bacterial biofilms [54].

In conclusion, this research advances the field of engineered living materials by providing a systematic understanding of how synthetic adhesins can be used to tune rheological properties and by demonstrating the potential of rheology as a design tool for material structuring with applications ranging from bioprinting and tissue engineering to environmental remediation and smart material design. Integrating additional biological components, such as extracellular matrices, or exploring different bacterial strains might further expand the range of tunable properties. Applying these findings to eukaryotic systems could broaden their applicability in biomedical contexts [55, 56]. Additionally, developing computational models to predict ELM behavior based on molecular parameters could enhance the design effciency and scalability of these materials [2]. Finally, these interplays between cell-cell adhesion and external mechanical flow-stresses could also explain consortia morphology and patterning of natural microbial systems [54].

## Methods

### Plasmids and strains

The MG1655 *E. coli* strain obtained from *E. Coli* Genetic Stock Center (CGSC 6300) and S1 *E. coli* strain obtained from *E. Coli* Genetic Stock Center (CGSC 6338) were used for all experiments in this study. Plasmids were transformed into chemically competent cells following standard protocol [10]. Plasmids were sourced from our earlier work [10]:

pDSG287 (pSB4A3_TetR_pTet_Neae2v1_EPEA) (GenBank ID MH492379);

pDSG320 (pSB3K3_TetR_pTet_Neae2v1_antiEPEA) (GenBank ID MH492391);

pDSG288 (pSB4A3_TetR_pTet_Neae2v1_P53TA) (GenBank ID MH492378);

pDSG321 (pSB3K3_TetR_pTet_Neae2v1_antiP53TA) (GenBank ID MH492393).

### Cell growth

A preculture (3 mL) of homophilic p53TA ot heterophilic EPEA *E. coli* strain was grown overnight. Next day, the preculture was transferred to 500 mL LB supplemented with the corresponding antibiotics and inducer concentration. For the elongated cells, 90 minutes prior to cell pellet preparation, cephalexin was added to 60g/ml.

### Cell pellet preparation

24 hours after induction, the cells were harvested by centrifugation at 5100 rpm for 10 minutes and the supernatant was removed. We used a centrifugation speed similar to the work done by other research groups [17]. We mixed the complementary cell cultures and ensured that the proper ratios were used by measuring the O.D. 600.

### Rheology measurements

The mechanical properties of the cell pellets were assessed using the Discovery Hybrid Rheometer 2 (TA Instruments, New Castle, DE, USA) with a sandblasted 1^°^, 20 mm cone-plate geometry through small amplitude oscillatory shear rhe-ology. Prior to each experiment, inertia and friction of the geometry and rotational mapping calibrations were performed. All experiments were conducted at a constant temperature of 25^°^C, facilitated by a Peltier temperature-controlled stage. 200 µL of each sample was spread evenly throughout the stage before lowering the gap height to 636 µm. Strain sweeps were performed from 0.1 to 10 % shear strain at a constant angular frequency of 10 rad s^−1^. G’ and G” were calculated by averaging the values within the linear viscoelastic region (0.1 – 10 %). For determining the viscosity of the cell pellets, a flow sweep was performed by applying a shear rate over a range of 0.01 to 1000 1/s.

### Statistics and reproducibility

Experimental replicates are indicated throughout the text. Statistical analysis was performed using GraphPad Prism software, with a p-value of less than 0.05 considered statistically significant. Unpaired two-tailed Student’s t-tests were used for comparisons between two conditions. No data were excluded from the analyses.

## Data and code availability

All analyzed data is available in the manuscript or the supplemental materials. Raw data and modeling scripts are available upon reasonable request.

## Acknowledgements

We thank members of the Riedel-Kruse lab for stimulating feedback and discussions. We thank the Imaging Cores and the University of Arizona Research, Innovation & Impact (RII) o!ce and the Technology Initiative Fund/Improving Health (TRIF) initiative. Funding was provided by the College of Science and MCB Department at the University of Arizona, NSF grant 2214020, NSF grant 2229070, NIH grant GM145893 (I.H.R-K.). S.Y.K. and M.K. were supported by NSF CAREER Grant DMR-2143126.

## Author contributions

S.A.C. and K.H.: Conceptualization, Data Curation, Formal analysis, Investigation (all experiments), Methodology, Validation, Visualization, Writing - original draft, Writing - review & editing. S.Y.K. and C.C.: Data Curation, Formal analysis, Investigation (initial rheology experiments). M.K.: Methodology, Resources, Supervision. I.H.R-K.: Conceptualization, Formal Analysis, Funding Acquisition, Project Administration, Resources, Supervision, Writing - original draft, Writing - review & editing. All authors have read and approved the manuscript.

## References

[1] Charlie Gilbert and Tom Ellis. Biological engineered living materials: growing functional materials with genetically programmable properties. ACS synthetic biology, 8(1):1–15, 2018.

[2] Wil V Srubar. Engineered living materials: taxonomies and emerging trends. Trends in biotechnology, 39(6):574–583, 2021.

[3] Andrés Díaz Lantada, Jan G Korvink, and Monsur Islam. Taxonomy for engineered living materials. Cell Reports Physical Science, 3(4), 2022.

[4] Xiaofan Jin and Ingmar H Riedel-Kruse. Optogenetic patterning generates multi-strain biofilms with spatially distributed antibiotic resistance. Nature Communications, 15(1):9443, 2024.

[5] Xiaofan Jin and Ingmar H Riedel-Kruse. Biofilm lithography enables high-resolution cell patterning via optogenetic adhesin expression. Proceedings of the National Academy of Sciences, 115(14):3698–3703, 2018.

[6] Honesty Kim, Dominic J Skinner, David S Glass, Alexander E Hamby, Bradey AR Stuart, Jörn Dunkel, and Ingmar H Riedel-Kruse. 4-bit adhesion logic enables universal multicellular interface patterning. Nature, 608(7922):324–329, 2022.

[7] Rohit Ruhal and Rashmi Kataria. Biofilm patterns in gram-positive and gram-negative bacteria. Microbiological Research, 251:126829, 2021.

[8] Minghui Xiao, Shuyi Lv, and Chunlei Zhu. Bacterial patterning: A promising biofabrication technique. ACS Applied Bio Materials, 7(12):8008–8018, 2024.

[9] Içvara Barbier, Hadiastri Kusumawardhani, and Yolanda Schaerli. Engineering synthetic spatial patterns in microbial populations and communities. Current Opinion in Microbiology, 67:102149, 2022.

[10] David S Glass and Ingmar H Riedel-Kruse. A synthetic bacterial cell-cell adhesion toolbox for programming multi-cellular morphologies and patterns. Cell, 174(3):649–658, 2018.

[11] Stefana A Costan, Paul M Ryan, Honesty Kim, Charles W Wolgemuth, and Ingmar H Riedel-Kruse. Biophysical characterization of synthetic adhesins for predicting and tuning engineered living material properties. Matter, 7(6):2125–2143, 2024.

[12] Adam J Stevens, Andrew R Harris, Josiah Gerdts, Ki H Kim, Coralie Trentesaux, Jonathan T Ramirez, Wesley L McKeithan, Faranak Fattahi, Ophir D Klein, Daniel A Fletcher, et al. Programming multicellular assembly with synthetic cell adhesion molecules. Nature, 614(7946):144–152, 2023.

[13] George Chao, Timothy M Wannier, Clair Gutierrez, Nathaniel C Borders, Evan Appleton, Anjali Chadha, Tina Lebar, and George M Church. helixcam: A platform for programmable cellular assembly in bacteria and human cells. Cell, 185(19):3551–3567, 2022.

[14] Barry Canton, Anna Labno, and Drew Endy. Refinement and standardization of synthetic biological parts and devices. Nature biotechnology, 26(7):787–793, 2008.

[15] Honesty Kim, Xiaofan Jin, David S Glass, and Ingmar H Riedel-Kruse. Engineering and modeling of multicellular morphologies and patterns. Current opinion in genetics & development, 63:95–102, 2020.

[16] Tzu-Chieh Tang, Bolin An, Yuanyuan Huang, Sangita Vasikaran, Yanyi Wang, Xiaoyu Jiang, Timothy K Lu, and Chao Zhong. Materials design by synthetic biology. Nature Reviews Materials, 6(4):332–350, 2021.

[17] Baizhu Chen, Wei Kang, Jing Sun, Runtao Zhu, Yue Yu, Aiguo Xia, Mei Yu, Meng Wang, Jinyu Han, Yixuan Chen, et al. Programmable living assembly of materials by bacterial adhesion. Nature Chemical Biology, 18(3):289–294, 2022.

[18] Lina M González, Nikita Mukhitov, and Christopher A Voigt. Resilient living materials built by printing bacterial spores. Nature chemical biology, 16(2):126–133, 2020.

[19] See-Yeun Ting, Esteban Martínez-García., Shuo Huang, Savannah K Bertolli, Katherine A Kelly, Kevin J Cutler, Elizabeth D Su, Hui Zhi, Qing Tang, Matthew C Radey, et al. Targeted depletion of bacteria from mixed populations by programmable adhesion with antagonistic competitor cells. Cell host & microbe, 28(2):313–321, 2020.

[20] Kenneth Timmis, James Kenneth Timmis, Harald Brüssow, and Luis Ángel Fernández. Synthetic consortia of nanobody-coupled and formatted bacteria for prophylaxis and therapy interventions targeting microbiome dysbiosis-associated diseases and co-morbidities. Microbial biotechnology, 12(1):58–65, 2019.

[21] Thomas K Wood, Ilke Gurgan, Ethan T Howley, and Ingmar H Riedel-Kruse. Converting methane into electricity and higher-value chemicals at scale via anaerobic microbial fuel cells. Renewable and Sustainable Energy Reviews, 188:113749, 2023.

[22] Allen Y Chen, Zhengtao Deng, Amanda N Billings, Urartu OS Seker, Michelle Y Lu, Robert J Citorik, Bijan Zakeri, and Timothy K Lu. Synthesis and patterning of tunable multiscale materials with engineered cells. Nature materials, 13(5):515–523, 2014.

[23] Maayan Lufton, Or Bustan, Bat-hen Eylon, Ella Shtifman-Segal, Tsuf Croitoru-Sadger, Alona Shagan, Ayelet Shabtay-Orbach, Enav Corem-Salkmon, Judith Berman, Abraham Nyska, et al. Living bacteria in thermoresponsive gel for treating fungal infections. Advanced Functional Materials, 28(40):1801581, 2018.

[24] Jiaofang Huang, Suying Liu, Chen Zhang, Xinyu Wang, Jiahua Pu, Fang Ba, Shuai Xue, Haifeng Ye, Tianxin Zhao, Ke Li, et al. Programmable and printable bacillus subtilis biofilms as engineered living materials. Nature chemical biology, 15(1):34–41, 2019.

[25] Shrikrishnan Sankaran, Judith Becker, Christoph Wittmann, and Aránzazu Del Campo. Optoregulated drug release from an engineered living material: self-replenishing drug depots for long-term, light-regulated delivery. Small, 15(5):1804717, 2019.

[26] Liliang Ouyang, Rui Yao, Yu Zhao, and Wei Sun. E”ect of bioink properties on printability and cell viability for 3d bioplotting of embryonic stem cells. Biofabrication, 8(3):035020, 2016.

[27] Jana Stepanovska, Monika Supova, Karel Hanzalek, Antonin Broz, and Roman Matejka. Collagen bioinks for bio-printing: a systematic review of hydrogel properties, bioprinting parameters, protocols, and bioprinted structure characteristics. Biomedicines, 9(9):1137, 2021.

[28] Yanyi Wang, Qian Zhang, Changhao Ge, Bolin An, and Chao Zhong. Programmable bacterial biofilms as engineered living materials. Accounts of Materials Research, 5(7):797–808, 2024.

[29] Rolf Lutz and Hermann Bujard. Independent and tight regulation of transcriptional units in escherichia coli via the lacr/o, the tetr/o and arac/i1-i2 regulatory elements. Nucleic acids research, 25(6):1203–1210, 1997.

[30] Mare Whitlow, Brian A Bell, Sheau-Line Feng, David Filpula, Karl D Hardman, Steven L Hubert, Michele L Rollence, James F Wood, Margaret E Schott, Diane E Milenic, et al. An improved linker for single-chain fv with reduced aggregation and enhanced proteolytic stability. Protein Engineering, Design and Selection, 6(8):989–995, 1993.

[31] Tony Y-C Tsai, Mateusz Sikora, Peng Xia, Tugba Colak-Champollion, Holger Knaut, Carl-Philipp Heisenberg, and Sean G Megason. An adhesion code ensures robust pattern formation during tissue morphogenesis. Science, 370(6512):113–116, 2020.

[32] Allen P Liu, Eric A Appel, Paul D Ashby, Brendon M Baker, Elisa Franco, Luo Gu, Karmella Haynes, Neel S Joshi, April M Kloxin, Paul HJ Kouwer, et al. The living interface between synthetic biology and biomaterial design. Nature materials, 21(4):390–397, 2022.

[33] Claude Verdier. Rheological properties of living materials. from cells to tissues. Computational and Mathematical Methods in Medicine, 5(2):67–91, 2003.

[34] M Keentok and RI Tanner. Cone-plate and parallel plate rheometry of some polymer solutions. Journal of Rheology, 26(3):301–311, 1982.

[35] Thomas Mezger. The rheology handbook: for users of rotational and oscillatory rheometers. European Coatings, 2020.

[36] A Franck and TI Germany. Viscoelasticity and dynamic mechanical testing. TA Instruments, New Castle, DE, USA AN004, 1993.

[37] Yankai Liu, Qingsong Zhang, and Rentai Liu. E”ect of particle size distribution and shear rate on relative viscosity of concentrated suspensions. Rheologica Acta, 60:763–774, 2021.

[38] Dong Yong Park and Seong Jin Park. Particle size-dependent viscosity behavior of a suspension using image processing. Powder Technology, 339:686–694, 2018.

[39] Willi Pabst, Eva Gregorová, and Christoph Berthold. Particle shape and suspension rheology of short-fiber systems. Journal of the European Ceramic Society, 26(1-2):149–160, 2006.

[40] Zhuoqing An, Yanling Zhang, Qi Li, Haoran Wang, Zhancheng Guo, and Jesse Zhu. E”ect of particle shape on the apparent viscosity of liquid–solid suspensions. Powder technology, 328:199–206, 2018.

[41] Laura K Rivera-Tarazona, Tarjani Shukla, Kanwar Abhay Singh, Akhilesh K Gaharwar, Zachary T Campbell, and Taylor H Ware. 4d printing of engineered living materials. Advanced Functional Materials, 32(4):2106843, 2022.

[42] Alison E Patteson, Arvind Gopinath, and Paulo E Arratia. Active colloids in complex fluids. Current Opinion in Colloid & Interface Science, 21:86–96, 2016.

[43] David S Glass and Uri Alon. Programming cells and tissues. Science, 361(6408):1199–1200, 2018.

[44] Christophe Ancey. Role of lubricated contacts in concentrated polydisperse suspensions. Journal of Rheology, 45(6):1421–1439, 2001.

[45] René Van der Ploeg, Jolanda Verheul, Norbert OE Vischer, Svetlana Alexeeva, Eelco Hoogendoorn, Marten Postma, Manuel Banzhaf, Waldemar Vollmer, and Tanneke Den Blaauwen. Colocalization and interaction between elongasome and divisome during a preparative cell division phase in e scherichia coli. Molecular microbiology, 87(5):1074–1087, 2013.

[46] Rajinder Pal. Complex shear modulus of concentrated suspensions of solid spherical particles. Journal of colloid and interface science, 245(1):171–177, 2002.

[47] Jan KG Dhont and Willem J Briels. Viscoelasticity of suspensions of long, rigid rods. Colloids and surfaces A: Physicochemical and engineering aspects, 213(2-3):131–156, 2003.

[48] M Mohseni and DG Allen. The e”ect of particle morphology and concentration on the directly measured yield stress in filamentous suspensions. Biotechnology and bioengineering, 48(3):257–265, 1995.

[49] Christophe Ancey and Hélene Jorrot. Yield stress for particle suspensions within a clay dispersion. Journal of Rheology, 45(2):297–319, 2001.

[50] Michael Florea, Henrik Hagemann, Gabriella Santosa, James Abbott, Chris N Micklem, Xenia Spencer-Milnes, Laura de Arroyo Garcia, Despoina Paschou, Christopher Lazenbatt, Deze Kong, et al. Engineering control of bacterial cellulose production using a genetic toolkit and a new cellulose-producing strain. Proceedings of the National Academy of Sciences, 113(24):E3431–E3440, 2016.

[51] M Tarek Abdelwahab, Ebuzer Kalyoncu, Tugce Onur, MehmetZ Baykara, and Urartu Ozgur Safak Seker. Genetically-tunable mechanical properties of bacterial functional amyloid nanofibers. Langmuir, 33(17):4337–4345, 2017.

[52] Howard Brenner. Rheology of a dilute suspension of axisymmetric brownian particles. International journal of multiphase flow, 1(2):195–341, 1974.

[53] Yujia Jiang, Ruofan Wu, Wenming Zhang, Fengxue Xin, and Min Jiang. Construction of stable microbial consortia for e”ective biochemical synthesis. Trends in Biotechnology, 41(11):1430–1441, 2023.

[54] Federica Recupido, Giuseppe Toscano, Rosarita Tatè, Maria Petala, Sergio Caserta, Thodoris D Karapantsios, and Stefano Guido. The role of flow in bacterial biofilm morphology and wetting properties. Colloids and Surfaces B: Biointerfaces, 192:111047, 2020.

[55] Veronika Magdanz, Samuel Sanchez, and Oliver G Schmidt. Development of a sperm-flagella driven micro-bio-robot. Advanced materials, 25(45):6581–6588, 2013.

[56] Caroline Cvetkovic, Ritu Raman, Vincent Chan, Brian J Williams, Madeline Tolish, Piyush Bajaj, Mahmut Selman Sakar, H Harry Asada, M Taher A Saif, and Rashid Bashir. Three-dimensionally printed biological machines powered by skeletal muscle. Proceedings of the National Academy of Sciences, 111(28):10125–10130, 2014.

