## Supplemental text for "Tuning viscoelasticity and fine structure of living materials via synthetic adhesion logic and rheological perturbations"

### Supplementary information

#### Supplementary Note 1: Introduction of synthetic adhesion (dual-component) in ELMs

We compared ELMs consisting of wild-type (wt) or uninduced heterophilic Nb3/Ag3 cells. We measured their viscosity through flow sweep tests and their viscoelasticity through strain amplitude sweep (Fig. S1). We determined that there is no statistical difference between these constructs (Students's t-test, $p \geq 0.1$). Therefore, we can disregard the possibility of leaky expression and combine the data.

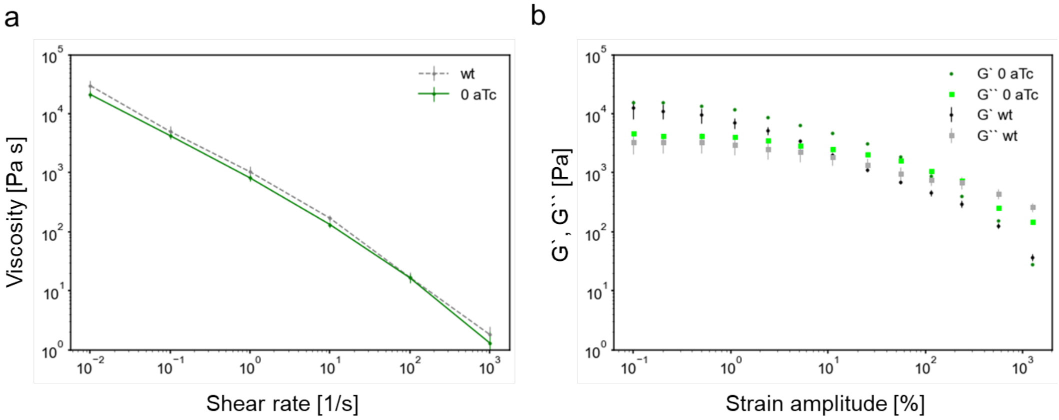


**Fig. S 1: Comparison between the rheological properties of the material consisting of wt and uninduced heterophilic cells.** **a** Measured viscosity for the wt and the uninduced (0 aTc) heterophilic Nb3-Ag3 cells. **b** Measured storage (G′) and loss (G″) moduli for the wt and the uninduced (0 aTc) heterophilic Nb3/Ag3 cells. Data presented as mean ± sem, unless noted otherwise.

#### Supplementary Note 2: Material tuning via the adhesin strength

We imaged the heterophilic Nb3/Ag3 cell mixtures before centrifugation and we observed in the confocal images that the cluster sizes increased with the inducer concentration (Fig. S2).

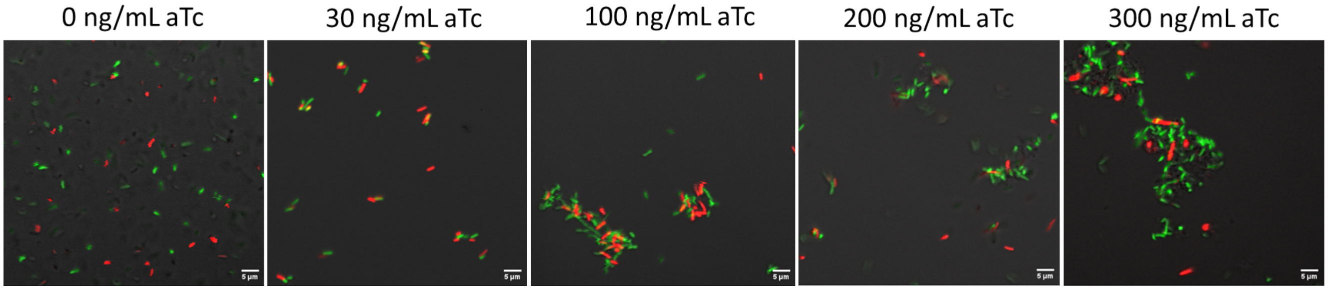


**Fig. S 2: Confocal images of the heterophilic Nb3-Ag3 cell mixtures under different leveles of induction in solution**

We then measured the viscosity of the ELMs under different levels of induction (Fig. S3a) and consequently, adhesin strength ($n = 4$ repeats across 2 days except for 0 and 100 ng/mL aTc at 0.1, 1, 10, 100, 1000 $s^{-1}$ shear rates where there are 7 and 9 repeats across 4 and 5 days, respectively). At higher shear rates (310 $s^{-1}$) we observed a gradual increase in the viscosity with the inducer concentration, whereas at lower shear rates (0.1 $s^{-1}$), the extreme levels of induction presented higher viscosity than the intermediary levels. We next quantified the dependence of the viscosity on the induction level. We determined the inducer concentration that produces half of the maximal viscosity ($C_{1/2}$) and the Hill coefficient (m) for high shear rates (310 $s^{-1}$) (Main text). Similarly, we fitted the Hill function for the intermediary shear rate level (3 $s^{-1}$), resulting in a $C_{1/2}$ and m not statistically different from 0 (Fig. S3b). For the low shear rate (0.1 $s^{-1}$) we were unable to fit any meaningful Hill curve (Fig. S3c).

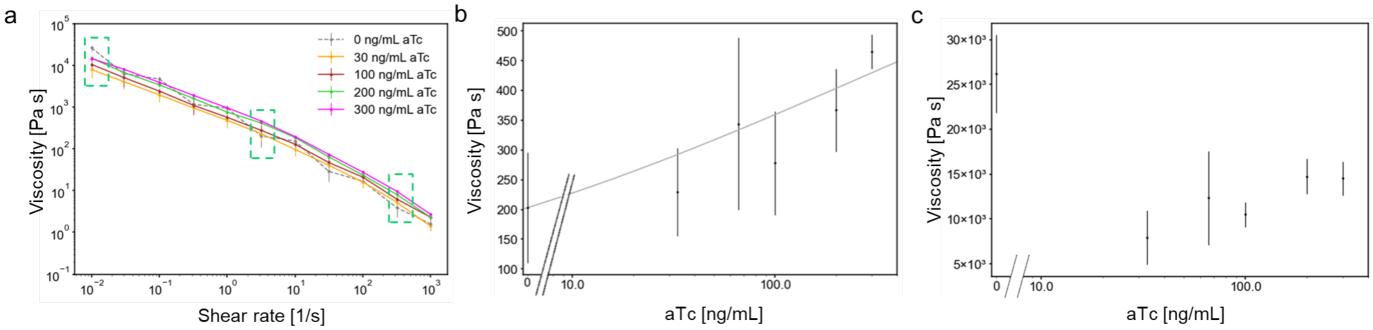


**Fig. S 3: The dependence of the ELM viscosity on the adhesin strength.** **a** Measured viscosity for materials under various levels of induction (0 - 300 ng/mL aTc) for a shear rate interval between 0.1 and 1000 $s^{-1}$. Dotted boxes represent the shear rate values chosen to investigate the Hill response of the viscosity on the inducer concentration. **b** Measured viscosity for materials under various levels of induction (0 - 300 ng/mL aTc) at a constant shear rate of 3 $s^{-1}$. Solid line represents the fitted Hill function. **b** Measured viscosity for materials under various levels of induction (0 - 300 ng/mL aTc) at a constant shear rate of 0.1 $s^{-1}$.

We were interested in how the material structure varies during the rheology measurements, therefore we took samples from the ELM at different induction levels before rheology, after one measurement and after going backwards with the shear rate. We dissolved the sample in PBS and we imaged the clusters (Fig. S4a,b,c). We determined the cluster size distribution for all three time points for 100 ng/mL aTc (Fig. 2c) and for 300 ng/mL aTc (Fig. S4d). At maximal induction before rheology the cluster sizes were 109 ± 35, whereas after rheology they remained constant at 63 ± 28.

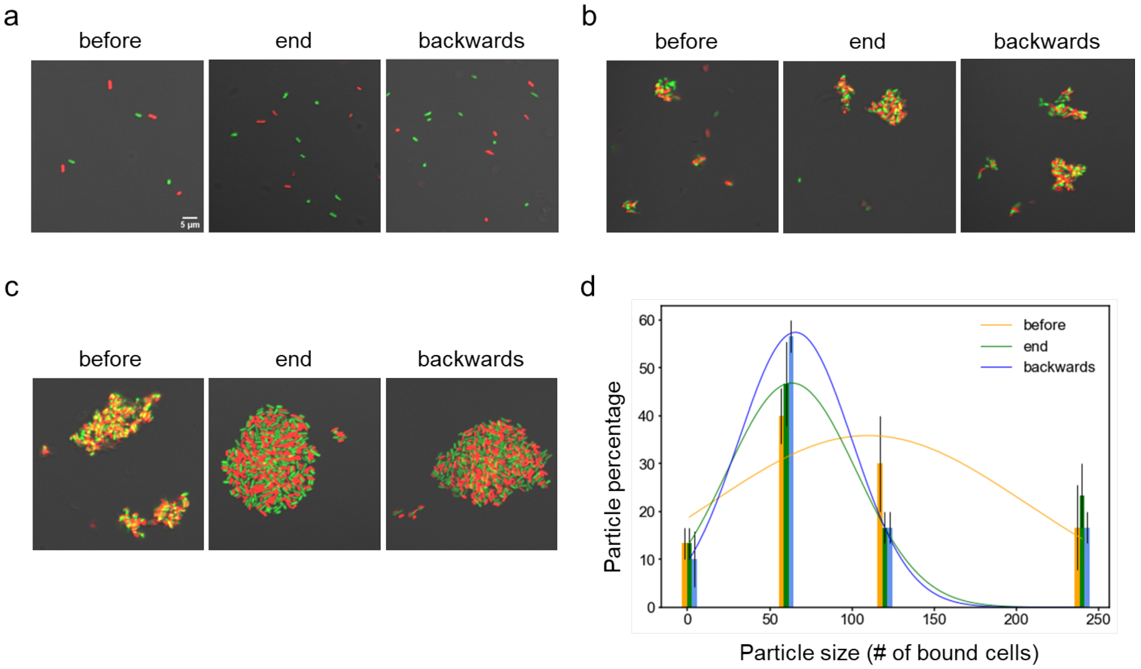


**Fig. S 4: Material microscopic structure changes during rheology.** Confocal images of the cell clusters inside the ELM sampled before, at the end on one measurement and after an additional backwards measurement for **a** 0 ng/mL aTc, **b** 100 ng/mL aTc, **c** 300 ng/mL aTc. **d** Cluster size distribution for the Nb3-Ag3 material dissolved in solution before viscosity measurement (orange), at the end of the measurement (green) and after going backwards with the shear rate (blue) for maximal induction.

We measured the loss and storage modulus for different induction levels. We then determined the yield point (yield stress) which is the value of the shear stress at the limit of the LVE region and the flow point (flow stress) which is the value of the shear stress at the crossover point $G' = G''$ (Main text).

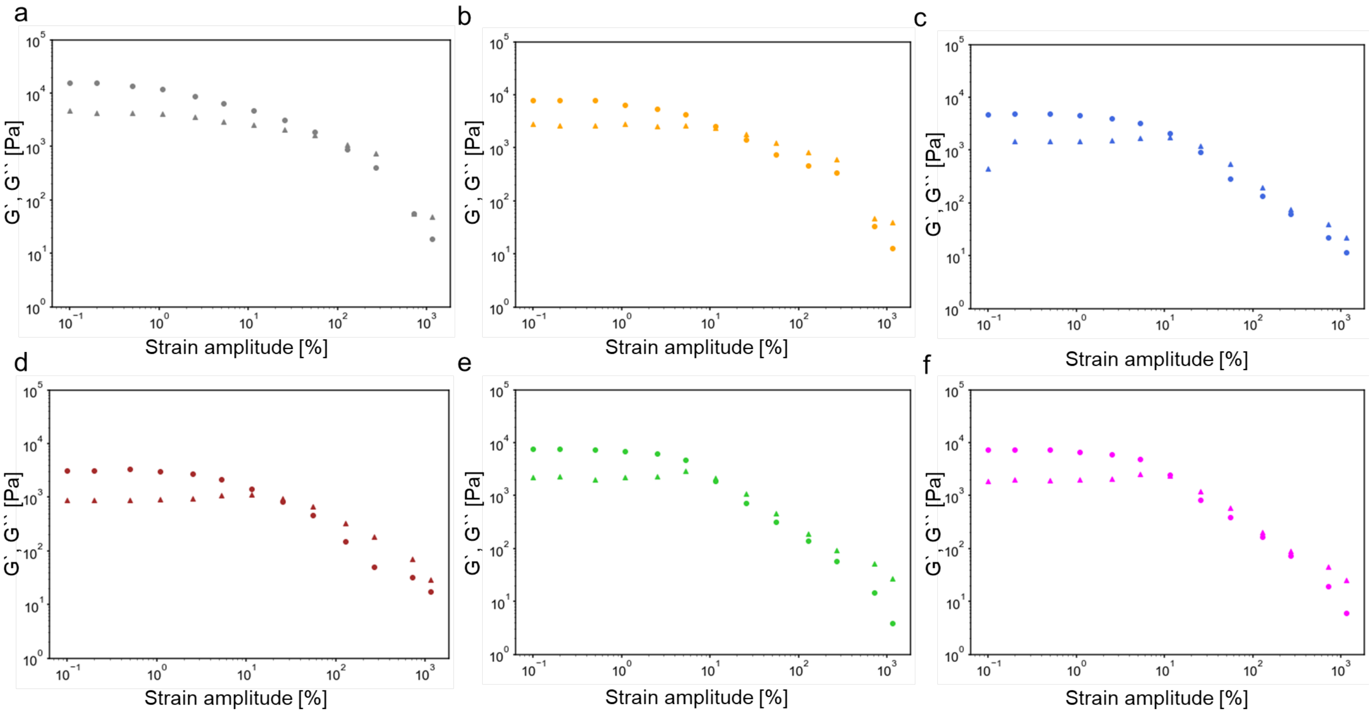


**Fig. S 5: Measured material viscoelasticity for different induction levels.** Storage (circles) and loss (triangles) modulus measured for **a** 0 ng/mL aTc **b** 30 ng/mL aTc, **c** 66 ng/mL aTc, **d** 100 ng/mL aTc, **e** 200 ng/mL aTc, **f** 300 ng/mL aTc.

#### Supplementary Note 3: Material tuning via the cell shape

We investigated the viscoelasticity of the ELMs consisting of different cell shapes that were either induced or not.

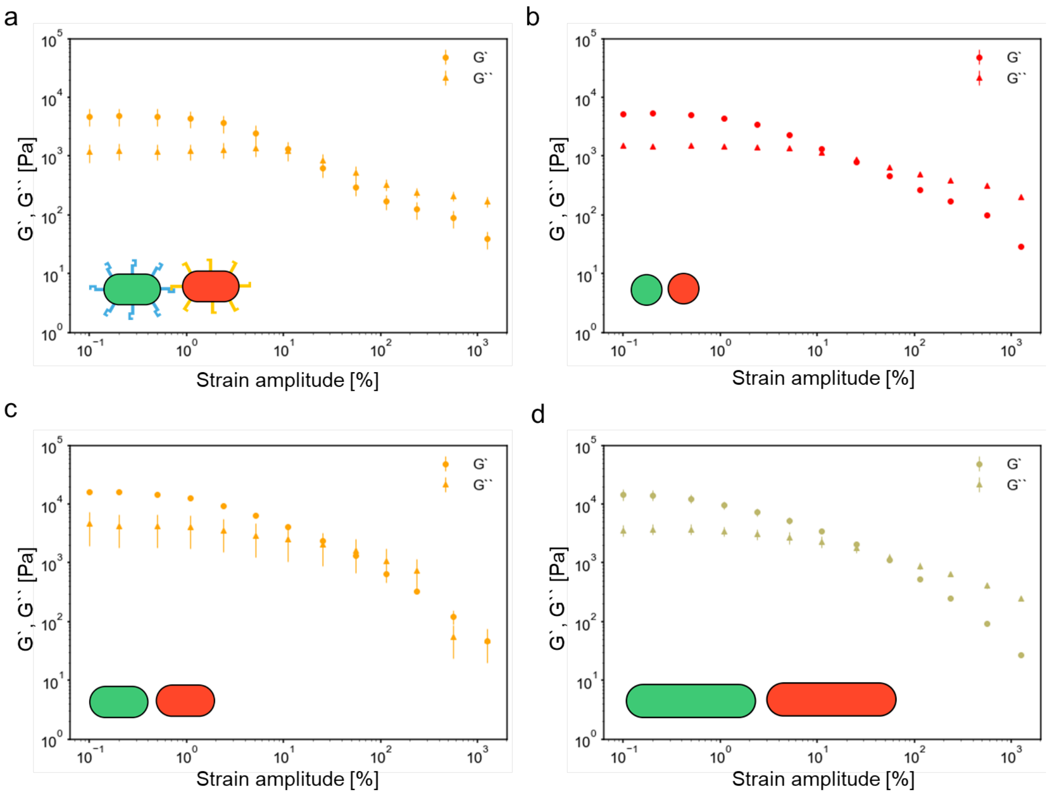


**Fig. S 6: Viscoelasticity of different constructs.** **a** Rod-shaped cells induced with 100 ng/mL aTc ($n = 4$ repeats across 2 days), **b** uninduced spherical cells ($n = 2$ repeats across 1 day), **c** uninduced rod-shaped cells ($n = 3$ repeats across 2 days), **c** uninduced elongated cells ($n = 2$ repeats across 1 day).

### Supplementary Note 4: Material tuning via higher-level adhesion (tri-component) and rheology

We repeated the experiment from Fig. 5 f-i by keeping the same mixing ratio between the red and the blue cells but without adding the filler cells - Nb3 (Mixture 4). We observed that after rheology the cluster size was similar to using induced filler cells but not as increased as it was by using uninduced filler cells. This was done by counting the number of red cells per cluster after the material was dissolved in PBS after rheology measurements.

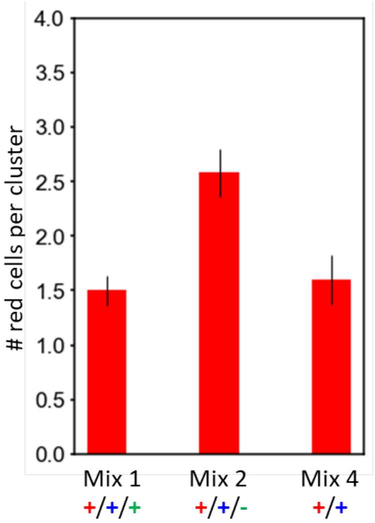


**Fig. S 7: Comparison for the number of red cels per cluster in the cell suspension after rheology between Mixtures 1, 2 and 4.**
